# White Matter Alterations in Young Children with Prenatal Alcohol Exposure

**DOI:** 10.1101/2021.01.05.425489

**Authors:** Preeti Kar, Jess E. Reynolds, Melody N. Grohs, W. Ben Gibbard, Carly McMorris, Christina Tortorelli, Catherine Lebel

## Abstract

Prenatal alcohol exposure (PAE) can lead to cognitive, behavioural, and social-emotional challenges. Previous neuroimaging research has identified alterations to brain structure in newborns, older children, adolescents, and adults with PAE; however, little is known about brain structure in young children. Extensive brain development takes place during early childhood; therefore, understanding the neurological profiles of young children with PAE is critical for early identification and effective intervention. We studied 54 children (5.21±1.11 years; 27 males) with confirmed PAE compared to 54 age- and sex-matched children without PAE. Children underwent diffusion tensor imaging between 2 and 7 years of age. Mean fractional anisotropy (FA) and mean diffusivity (MD) were obtained for 10 major white matter tracts, along with tract volume, axial and radial diffusivity (AD, RD). A univariate analysis of covariance was conducted to test for group differences (PAE vs. control) controlling for age, sex and tract volume. Our results reveal white matter microstructural differences between young children with PAE and unexposed controls. The PAE group had higher FA and/or lower MD (as well as lower AD and RD) in the genu and the body of the corpus callosum, as well as the bilateral uncinate fasciculus and pyramidal tracts. Our findings align with studies of newborns with PAE finding lower AD, but contrast those in older populations with PAE, which consistently report lower FA and higher MD. These findings may reflect premature development of white matter that may then plateau too early, leading to the lower FA/higher MD observed at older ages.

## Introduction

Prenatal alcohol exposure (PAE) affects the developing brain by altering neuronal and glial proliferation and migration, as well as disrupting synaptogenesis, gene transcription, and apoptotic processes (Goodlett et al., 2005; Wilhelm et al., 2016). Fetal alcohol spectrum disorder (FASD) is the neurodevelopmental disorder caused by PAE (Cook et al., 2015), and has an estimated prevalence of 2-5% in North America (May et al., 2014; Popova et al., 2019), considerably higher than most other neurodevelopmental disorders (Flannigan et al., 2018).

Cognitive, behavioural and social-emotional challenges are commonly observed in individuals with PAE and/or FASD, including lower cognitive and academic achievement (Mattson et al., 2019), impairments in expressive and receptive language (Wyper et al., 2011), motor deficits (Lucas et al., 2016), as well as a significantly increased risk of mental health disorders (Pei et al., 2011). Investigating the brain structure underlying these impairments can shed light upon the neurobiological effects of PAE teratogenesis.

Magnetic resonance imaging (MRI) has been used to study brain structure in newborns, school-aged children, adolescents, and adults with PAE, demonstrating widespread alterations such as reduced total brain volume and alterations to cortical and subcortical regional volumes, cortical thickness, and functional activation (Donald et al., 2015a; Nguyen et al., 2017). White matter appears to be particularly affected by PAE, with volume studies suggesting disproportionate reductions to white matter (Archibald et al., 2001; Jacobson et al., 2017; Nardelli et al., 2011) and diffusion tensor imaging (DTI) studies reporting extensive alterations to microstructure (Donald et al., 2015a; Ghazi Sherbaf et al., 2018; Lebel et al., 2008a). Measures of white matter microstructure, such as fractional anisotropy (FA) and mean, axial and radial diffusivity (MD, AD, RD), reflect properties such as myelination, axon diameter, and/or axon packing density. In small studies of newborns, PAE has been associated with lower AD in transcallosal, projection, and association fibers (Taylor et al., 2016) as well as lower AD in the right superior longitudinal fasciculus (Donald et al., 2015b). Furthermore, altered cerebellar white matter in newborns with PAE has been associated with neurobehavioral outcomes (Donald et al., 2015b). In school-aged children, adolescents and adults with PAE/FASD, lower FA and/or higher MD have been consistently noted throughout the brain (Donald et al., 2015a; Ghazi Sherbaf et al., 2018; Lebel et al., 2008a). Some studies have associated lower FA and/or higher MD with difficulties in processing speed (Fan et al., 2016), reading (Treit et al., 2013), working memory (Wozniak et al., 2009), mathematics (Lebel et al., 2010), and executive functioning (Treit et al., 2017), as well as eyeblink conditioning (Fan et al., 2015). Children with PAE also demonstrate altered development trajectories of white matter, with one study showing steeper decreases in MD in the inferior and superior longitudinal fasciculus and superior fronto-occipital fasciculus from childhood to adolescence (Treit et al., 2013).

There remains a significant gap in the literature on brain abnormalities in young children with PAE, which hinders the development of effective early assessment and intervention (Cook et al., 2015). Early childhood (∼2-7 years) is a period of extensive brain development, with rapid cognitive, behavioural, and social-emotional changes (Deoni et al., 2012; Hermoye et al., 2006; Pfefferbaum et al., 1994), as well as a period when many developmental delays first become apparent (Bitsko et al., 2016; Christensen et al., 2016; Visser et al., 2015). Identifying challenges as early as possible can facilitate early intervention, yet FASD is often not diagnosed until age 7 or older in many clinics (Cook et al., 2015). Here, we aimed to characterize white matter microstructure in young children with PAE using DTI. Based on previous neuroimaging literature in older populations with PAE and/or FASD (Lebel et al., 2008a; Treit et al., 2013), we hypothesized that young children with PAE would demonstrate lower FA and higher MD than unexposed controls across brain white matter tracts.

## Materials and Methods

### Participants

We recruited 57 children with confirmed PAE through caregiver support groups, early intervention services, and Alberta Children’s Services in Alberta, Canada. Exclusion criteria were birth before 34 weeks’ gestation, children for whom English was not a primary language, history of head trauma, a diagnosis of autism, cerebral palsy, epilepsy or any other medical or genetic disorder associated with serious motor or cognitive disability, and contraindications to MRI (e.g., metal implants, dental braces). Children with attention deficit hyperactivity disorder, learning disabilities, language delays, and/or mental health diagnoses were included, as these frequently co-occur with PAE. None of the children were diagnosed with FASD given that most clinics in Alberta do not assess children for FASD until they are at least 7-years-old. Of the 57 children recruited, 1 child was excluded for an incidental finding on the MRI scan and 2 children did not feel comfortable receiving an MRI scan, leaving 54 children with PAE with usable MRI data, aged 2.49 – 6.97 years (5.21±1.11 years; 27M/27F). The sample included one pair of twins, one pair of non-twin full-siblings, a group of 3 non-twin full-siblings, and 4 pairs of non-twin half-siblings.

Age- and sex-matched unexposed children were selected from an ongoing study of typical brain development (Reynolds et al., 2020; Reynolds et al., 2019). All control participants were born >37 weeks’ gestation, spoke English as a primary language, and had no contraindications to MRI scans as well as had no history of developmental disorders or brain trauma. These unexposed children were aged 2.49-6.97 years (5.21±1.11 years; 27M/27F). Parent/guardian written informed consent and child verbal assent were obtained for each subject. The University of Calgary Conjoint Health Research Ethics Board (CHREB) approved this study (REB14-2266, REB13-020).

### Assessment of Prenatal/Postnatal Adverse Exposures

For children with PAE, information pertaining to prenatal and postnatal exposures was obtained from each participant’s child welfare file (which contained information from birth families, social workers, police records, and medical files) and/or from semi-structured interviews with current caregivers, caseworkers, and/or birth families. Prenatal and postnatal profiles were evaluated according to our previously reported framework (Lebel et al., 2019). Among participants with PAE, 31% (n=17) had confirmed PAE greater than or equal to the threshold indicated by the Canadian Guidelines for Diagnosing FASD (Cook et al., 2015): ≥7 drinks in 1 week and/or 2 or more binge episodes (≥4 drinks at one time) during pregnancy; 69% (n=37) of participants had confirmed PAE or a lower or unspecified amount. 94% (51) children with PAE also had prenatal exposure to other substances; 46% (n=25) were exposed to cannabis, 33% (n=18) to tobacco, and 63% (n=34) to illicit substances such as stimulants, methamphetamines, or opioids. 74% (n=40) of participants with PAE had adverse postnatal experiences such as neglect, physical/sexual/emotional abuse, witnessing violence and/or substance use, and/or multiple caregiver transitions. The remaining 26% (n=14) of participants with PAE had no postnatal adverse exposures. The average age of stable placement, after which point there were no more postnatal adverse experiences (as defined above), ranged from 0 to 4.08 years (0.92±1.14 years). Control participants had confirmed absence of PAE and prenatal exposure to other substances based on prospective questionnaires and interviews completed with the mother during pregnancy, no reports of postnatal adversities (i.e., abuse, neglect), and were still residing with their biological parent(s) at the time of their MRI scan.

### Neuroimaging Data Acquisition

Children underwent an MRI scan at the Alberta Children’s Hospital on the research-dedicated GE 3T MR750w system with a 32-channel head coil. Children were not sedated for scanning, but families were given reading materials to prepare children at home and offered one or more practice sessions in an MRI simulator (Thieba et al., 2018). Foam padding was used to minimize head motion, and headphones with a projector and screen allowed children to watch a movie throughout the scan. Whole-brain diffusion weighted images were acquired in 4:03 minutes using single shot spin echo echo-planar imaging sequence with: 1.6 x 1.6 x 2.2 mm resolution (resampled to 0.78 x 0.78 x 2.2mm on scanner), TR = 6750 ms; TE = 79 ms, 30 gradient encoding directions at b=750 s/mm^2^, and 5 interleaved images without diffusion encoding at b=0 s/mm^2^.

### Neuroimaging Data Processing

DTI data was visually inspected for quality, and volumes with artifacts or motion corruption were removed according to our previous methods (Reynolds et al., 2019; Walton et al., 2018). Children with fewer than 18 diffusion weighted volumes and 2 non-diffusion weighted volume were eliminated (Reynolds et al., 2019; Walton et al., 2018). In this sample, unexposed controls had between 19-30 diffusion weighted volumes (mean±SD 28±3) and 3-5 non-diffusion weighted volumes (5±0) remaining while children with PAE had 18-30 diffusion weighted volumes (26±4) and 3-5 non-diffusion weighted volumes (5±1) remaining. There were no significant differences between groups for number of volumes. All data was preprocessed using ExploreDTI (V4.8.6) which involved corrections for signal drift, Gibbs ringing, subject motion, and eddy current distortions (Leemans et al., 2009). Next, semi-automated deterministic streamline tractography was used to delineate 10 major white matter tracts: the corpus callosum (genu, body, splenium), and the fornix as well as the left and right cingulum, pyramidal tract, uncinate fasciculus, superior longitudinal fasciculus (SLF), inferior longitudinal fasciculus (ILF), and inferior fronto-occipital fasciculus (IFOF) (Lebel et al., 2012a; Lebel et al., 2008a; Reynolds et al., 2019). All region of interest semi-automated tractography guides can be found at: https://doi.org/10.6084/m9.figshare.7603271 (Reynolds et al., 2020). The minimum FA threshold was set to 0.20 to initiate and continue tracking, and the angle threshold was set to 30° to minimize spurious fibers (Lebel et al., 2008a; Reynolds et al., 2019). All tracts were manually quality checked and additional exclusion regions were drawn as required to remove spurious fibers. FA, MD, AD, RD, and tract volume values were calculated for every tract, separately for left and right hemisphere where relevant (i.e., all tracts except the corpus callosum and fornix). It is important to note that tract volume obtained using tractography is a measure of the volume of white matter in a fiber bundle that exceeds the chosen FA threshold (0.20) and does not necessarily represent the true white matter volume of the tract (Lebel et al., 2011).

### Statistical Analysis

Using SPSS (Version 25), a univariate analysis of covariance (ANCOVA) was used to test for group differences in tract volume (each tract separately); age and sex were included as covariates because they are related to brain structure in early childhood (Reynolds et al., 2019). An ANCOVA was then used to test for group differences in diffusion parameters, FA and MD separately, in each tract while controlling for age, sex, and tract volume. To further investigate tracts showing significant group differences in FA or MD, an ANCOVA was conducted to test for group differences in AD and RD separately while controlling for age, sex, and tract volume. Three supplementary analyses were conducted separately for the tracts with significant FA or MD group differences: the first, to account for motion by adding the number of remaining diffusion-weighted volumes for each dataset as a covariate; the second, to account for prenatal substance exposure by adding cannabis, tobacco, and illicit drug use each as a binary (exposed/unexposed) covariate, and the third to account for postnatal adverse exposures by adding age at stable placement as a covariate. MD, AD, and RD values were scaled by 1000 for analyses to bring them to a similar scale as other measures. False discovery rate (FDR) was used to correct for 48 multiple comparisons for post-hoc testing (3 tract parameters – tract volume, FA, MD – for each of 16 individual tracts), with significance set at *q* < 0.05.

## Results

### Fractional anisotropy

Children with PAE had significantly higher FA in the genu and the body of the corpus callosum, the left cingulum, and bilateral pyramidal tracts. In the fornix, children with PAE had significantly lower FA than unexposed controls. All findings, with the exception of the left cingulum, survived FDR correction for multiple comparisons and remained significant after accounting for motion (Table 1, Figure 1).

**Table 1.** Group differences (children with PAE vs. unexposed control) in diffusion measures for individual tracts, controlling for age at scan, sex, and tract volume.

|  | Fractional Anisotropy |  | Mean Diffusivity |  | Tract Volume |  |
| --- | --- | --- | --- | --- | --- | --- |
|  | F | *p* (*q*) | F | *p* (*q*) | F | *p* (*q*) |
| Genu of the Corpus Callosum | **13.4** | **<0.001 (0.019)** | **8.1** | **0.005 (0.032)** | **7.3** | **0.008 (0.036)** |
| Body of the Corpus Callosum | **7.9** | **0.006 (0.031)** | 1.3 | 0.259 (0.414) | **11.5** | **<0.001 (0.012)** |
| Splenium of the Corpus Callosum | 0.8 | 0.376 (0.531) | 0.9 | 0.343 (0.515) | 4.4 | 0.038 (0.101) |
| Fornix | **7.2** | **0.008 (0.036)** | 3.0E-3 | 0.958 (0.978) | 0.017 | 0.895 (0.955) |
| Left Cingulum | 4.5 | 0.037 (0.111) | 0.1 | 0.805 (0.878) | 4.5 | 0.037 (0.104) |
| Right Cingulum | 2.7 | 0.102 (0.223) | 0.6 | 0.437 (0.599) | 0.1 | 0.798 (0.891) |
| Left Inferior Fronto-Occipital Fasciculus | 2.2 | 0.142 (0.284) | 1.1 | 0.304 (0.471) | **12.4** | **<0.001 (0.010)** |
| Right Inferior Fronto-Occipital Fasciculus | 0.007 | 0.934 (0.975) | 3.9 | 0.051 (0.129) | **13.2** | **<0.001 (0.010)** |
| Left Inferior Longitudinal Fasciculus | 2.1 | 0.149 (0.286) | 0.4 | 0.509 (0.471) | 3.0 | 0.086 (0.197) |
| Right Inferior Longitudinal Fasciculus | 1.5 | 0.226 (0.374) | 0.9 | 0.350 (0.509) | 0.1 | 0.757 (0.865) |
| Left Superior Longitudinal Fasciculus | 0.4 | 0.542 (0.685) | 0.2 | 0.628 (0.754) | 5.6 | 0.020 (0.069) |
| Right Superior Longitudinal Fasciculus | 0.5 | 0.462 (0.616) | 4.6E-4 | 0.983 (0.983) | **8.1** | **0.005 (0.036)** |
| Left Pyramidal Tract | **7.6** | **0.007 (0.034)** | 1.7 | 0.195 (0.347) | 3.6 | 0.061 (0.146) |
| Right Pyramidal Tract | **10.9** | **0.001 (0.013)** | 0.3 | 0.603 (0.742) | 5.0 | 0.028 (0.090) |
| Left Uncinate Fasciculus | 1.7 | 0.201 (0.345) | **6.9** | **0.010 (0.036)** | 2.6 | 0.110 (0.230) |
| Right Uncinate Fasciculus | 0.2 | 0.651 (0.762) | **8.9** | **0.004 (0.028)** | 2.0 | 0.162 (0.299) |

Findings that survive correction for multiple comparisons using false discovery rate are bolded.

**Figure 1.**
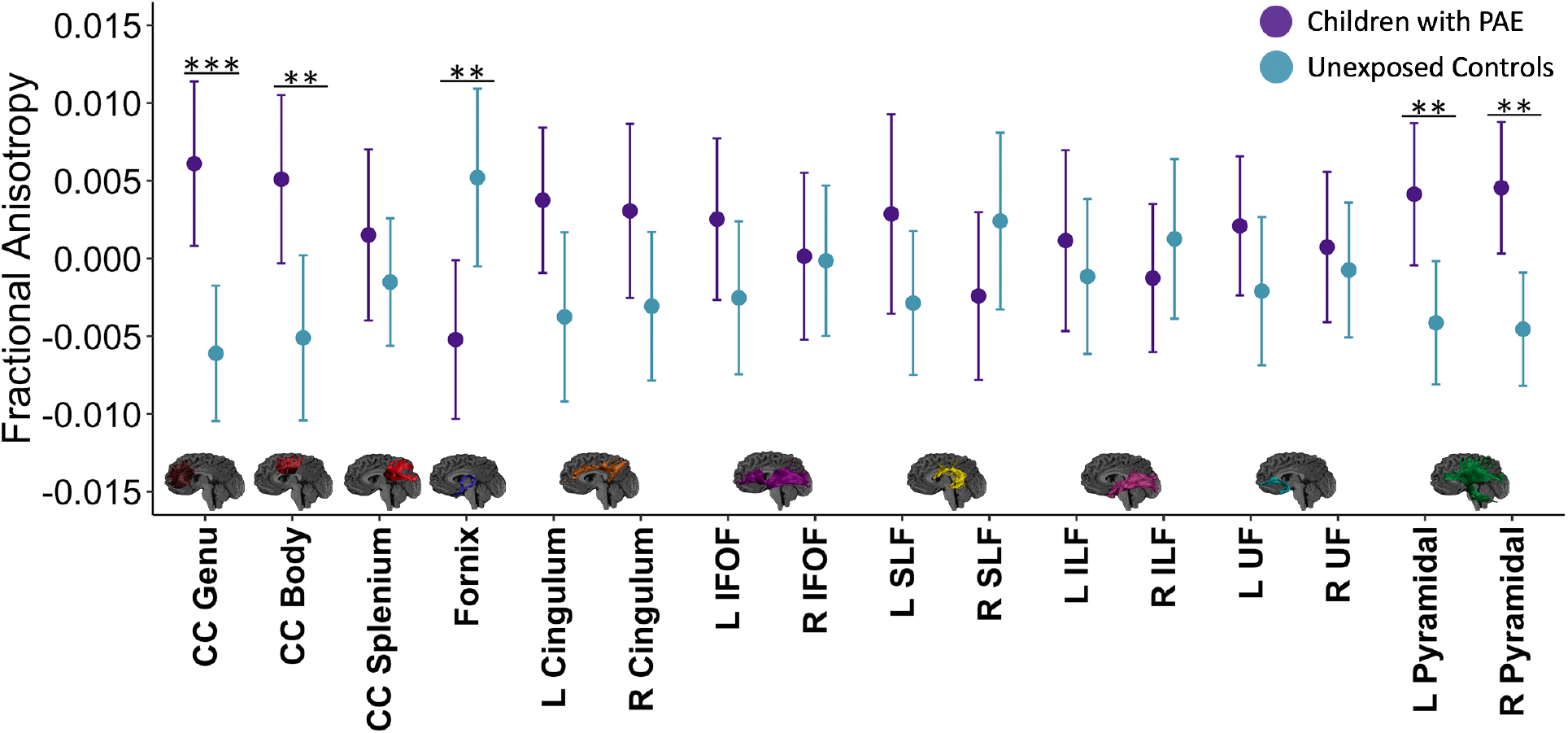
Group differences in FA. Children with PAE (purple) had significantly higher FA in genu and the body of the corpus callosum, and the bilateral pyramidal tracts compared to unexposed controls (blue). Children with PAE had significantly lower FA in the fornix compared to controls. Data is represented as mean FA ± 95% confidence interval for FA values for individual tracts. FA values are corrected for the participant’s age, sex, and tract volume. CC Genu = genu of the corpus callosum, CC Body = body of the corpus callosum, CC Splenium = splenium of the corpus callosum, UF = uncinate fasciculus. ***p<0.001, **p<0.01, *p<0.05.

### Mean diffusivity

Children with PAE had significantly lower MD in the genu of the corpus callosum and bilateral uncinate fasciculus compared to unexposed controls. All findings survived FDR correction for multiple comparisons and remained significant after accounting for motion (Table 1, Figure 2).

**Figure 2.**
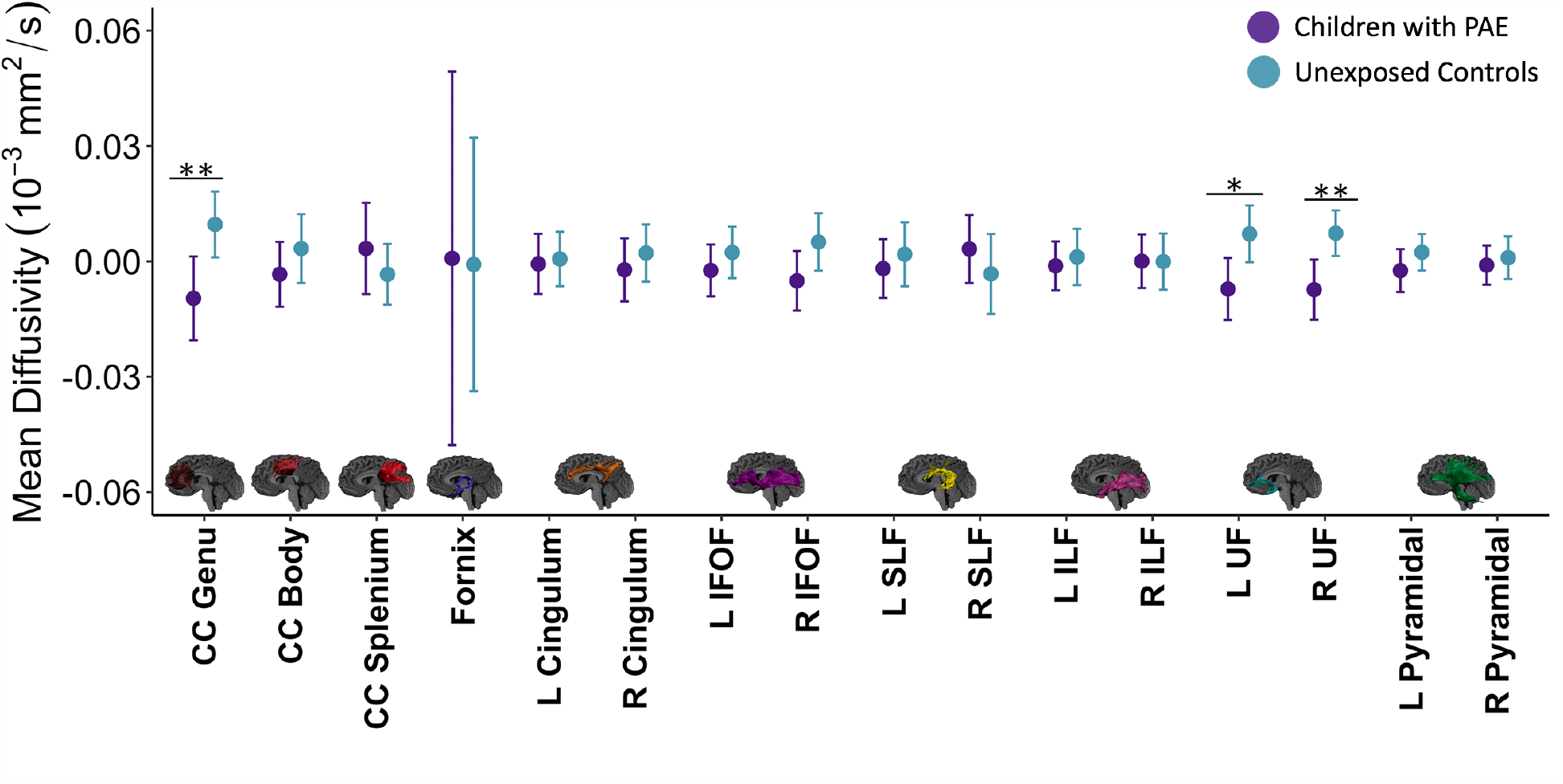
Group differences in MD. Children with PAE (purple) had significantly lower MD in the genu of the corpus callosum and the bilateral uncinate fasciculus compared to unexposed controls (blue). Data is represented as mean MD ± 95% confidence interval for MD values for individual tracts. MD values are corrected for the participant’s age, sex, and tract volume. CC Genu = genu of the corpus callosum, CC Body = body of the corpus callosum, CC Splenium = splenium of the corpus callosum, UF = uncinate fasciculus. ***p<0.001, **p<0.01, *p<0.05.

### Radial and axial diffusivity

Children with PAE had lower RD in the genu of the corpus callosum (*p* = 0.001) and the bilateral uncinate fasciculus (left: *p* = 0.013; right: *p* = 0.015) compared to unexposed controls. The PAE group showed lower AD in the left (*p* = 0.034) and the right (*p* = 0.006) uncinate fasciculus.

### Tract volume

Children with PAE had significantly lower tract volume in the corpus callosum (genu, body, and splenium), left cingulum, right SLF, right pyramidal tract, and bilateral IFOF compared to unexposed controls. The only tract with significantly higher volume in the PAE group was the left SLF. Findings in the body and genu of the corpus callosum, bilateral IFOF, and the right SLF survived FDR correction for multiple comparisons (Table 1, Figure 3).

**Figure 3.**
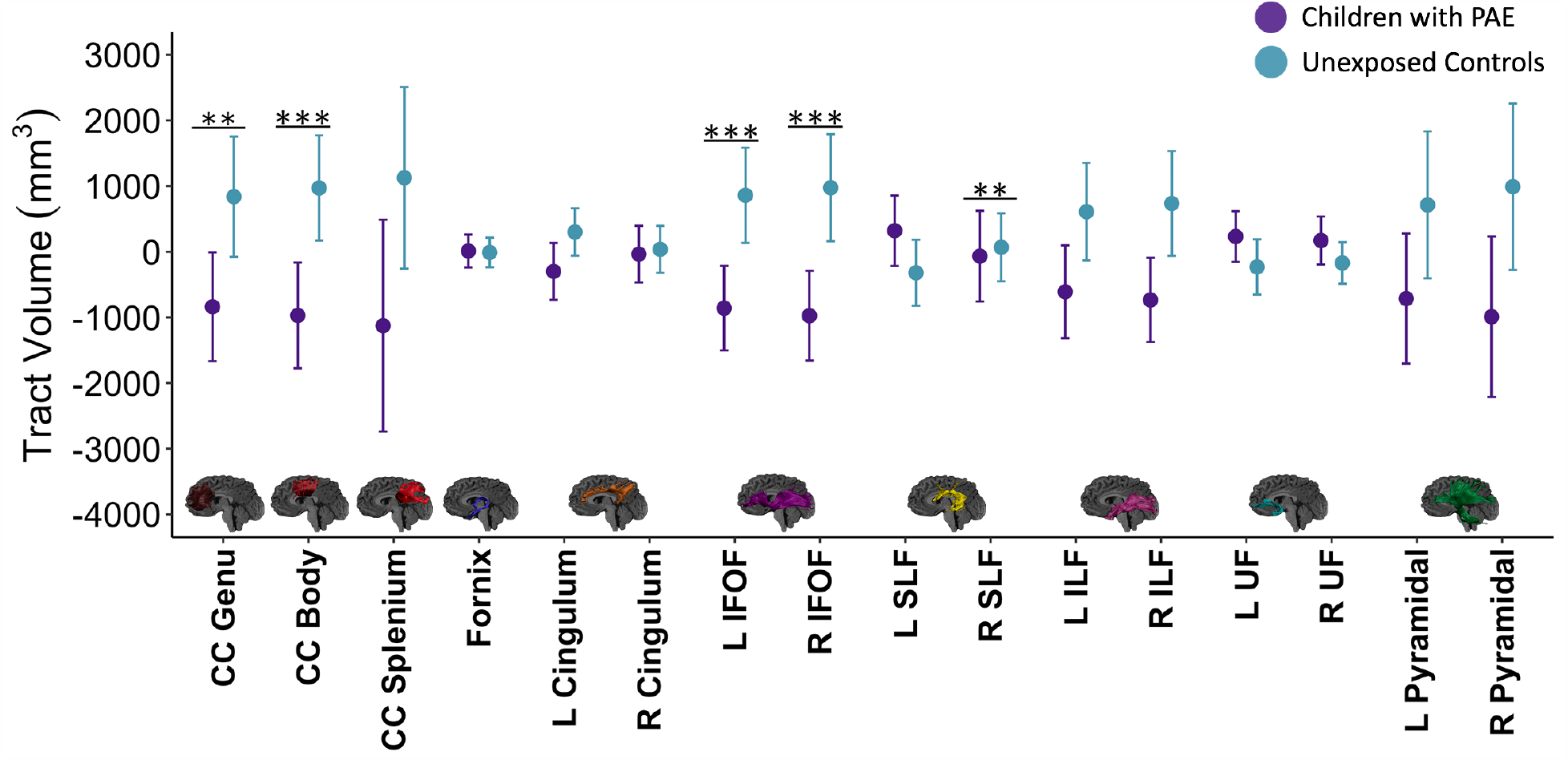
Group differences in tract volume. Children with PAE (purple) had significantly lower tract volume compared to unexposed controls in the genu and the body of the corpus callosum, as well as the right and left IFOF, and the right SLF. Data is represented as mean tract volume ± 95% confidence interval for tract volume values for individual tracts. Tract volume values are corrected for the participant’s age and sex. CC Genu = genu of the corpus callosum, CC Body = body of the corpus callosum, CC Splenium = splenium of the corpus callosum, UF = uncinate fasciculus. ***p<0.001, **p<0.01, *p<0.05.

### Other Prenatal and Postnatal Adverse Exposures

When accounting for other prenatal substance exposure (cannabis, tobacco, illicit drugs), group differences of FA in the body (p=0.004) and the genu (p=0.005) of the corpus callosum and bilateral pyramidal tracts (right: p=0.001; left: p=0.028) remained significant; the fornix no longer had significant group differences (p=0.67). MD group differences in the genu (p=0.021) of the corpus callosum and the right uncinate fasciculus (p=0.006) remained significant after controlling for other substance use; the left uncinate fasciculus group difference was no longer significant (p=0.19).

All group differences remained significant (p<0.05) after controlling for age at stable placement as a measure for postnatal adversities.

## Discussion

Here, we show altered white matter microstructure in young children with PAE for the first time. Higher FA and lower MD were noted in young children with PAE, which differs from previous findings in older children and adolescents (Donald et al., 2015a; Fan et al., 2016; Ghazi Sherbaf et al., 2018; Nguyen et al., 2017; Wozniak et al., 2011), though limited previous research in newborns has shown lower AD (Donald et al., 2015b; Taylor et al., 2016). Throughout childhood, FA increases and MD decreases with age across the brain (Lebel et al., 2011; Lebel et al., 2008b; Reynolds et al., 2019); thus, higher FA and lower MD may reflect premature development of white matter. Because previous studies consistently show lower FA and higher MD in older children and adolescents with PAE (Donald et al., 2015a; Ghazi Sherbaf et al., 2018; Nguyen et al., 2017), it is likely that premature development is then followed by early plateaus in developmental trajectories of white matter.

The corpus callosum facilitates interhemispheric communication (Aboitiz et al., 2003; Grohs et al., 2018) and is especially affected by PAE (Boronat et al., 2017; Yang et al., 2013). Previous research consistently reports higher FA and/or lower MD in the corpus callosum of individuals with PAE/FASD (Ma et al., 2005; Paolozza et al., 2017; Treit et al., 2013; Uban et al., 2017; Wozniak et al., 2009). Some isolated findings align more closely with our results, including lower genu MD (Lebel et al., 2008a) in children with FASD, higher FA in the body of the corpus callosum in boys with PAE (Uban et al., 2017), and lower AD in transcallosal pathways of 11 newborns with PAE (Taylor et al., 2016). The uncinate fasciculus, which connects frontal regions and limbic structures and is implicated in social-emotional processing and memory (Olson et al., 2015), and the pyramidal tracts, projection fibers implicated in motor performance (Jang, 2014; Yeo et al., 2014), also had higher FA in the PAE group. Previous research consistently reports lower FA and/or higher MD in the uncinate fasciculus and corticospinal tracts (Fan et al., 2016; Lebel et al., 2008a; O’Conaill et al., 2015; Paolozza et al., 2017; Treit et al., 2013; Uban et al., 2017) in older children and adolescents, though lower AD was reported in newborns (Taylor et al., 2016). These findings suggest that white matter differences associated with PAE are present across ages but may change directions between early and late childhood.

The only brain region that had lower FA in children with PAE was the fornix, a limbic tract related to memory (Ly et al., 2016). The fornix also undergoes rapid development during early childhood and reaches a developmental plateau earlier than other tracts (Lebel et al., 2012a; Reynolds et al., 2019). Thus, if white matter in children with PAE develops prematurely, the fornix would be the first to reverse from higher to lower FA compared to unexposed controls. The fornix also has high variability in diffusion parameters, in part due to its proximity to the ventricles, which may result in partial volume effects (Lebel et al., 2017). Finally, the FA differences in the fornix did not remain significant after controlling for other prenatal substance exposures, suggesting they may be less robust than the differences in the corpus callosum, uncinate, and pyramidal tracts.

Lower volume was noted across multiple white matter tracts, which aligns with extensive work showing smaller total brain volume (Donald et al., 2015a; Nguyen et al., 2017) and white matter volume (Archibald et al., 2001; Jacobson et al., 2017; Nardelli et al., 2011), related to PAE. Lower tract volume, coupled with higher FA and/or lower MD, may indicate that axons are more tightly packed together within the tract (Beaulieu, 2002). Both tract volume and FA increase with age, though they follow different trajectories (Lebel et al., 2012a; Moura et al., 2016; Reynolds et al., 2019), which may lead to different group profiles in older children with PAE (i.e., lower FA and lower volume) relative to unexposed controls.

This premature brain development suggested by our findings may reflect a compensatory neural mechanism caused by PAE where the brain tries to adapt to a challenging environment by achieving a mature structure sooner rather than following a prolonged development trajectory (Bick et al., 2016; Callaghan et al., 2016; Mehta et al., 2009; Whittle et al., 2013). Interestingly, premature development related to early postnatal adversity (e.g., institutionalization, maltreatment) has been noted most prominently within the limbic system, including the uncinate fasciculus (Gee et al., 2013; Mehta et al., 2009; Merz et al., 2018; Tottenham et al., 2010; Whittle et al., 2013). Deviation from typical brain development can alter windows of plasticity in populations with neurodevelopmental challenges (Fagiolini et al., 2011), and previous research has suggested altered brain plasticity in children with PAE, including earlier peaks for cortical volume (Lebel et al., 2012b) and steeper decreases in MD (Treit et al., 2013). Specific mechanisms that trigger the shift from brain overdevelopment to underdevelopment and alter brain plasticity require further investigation.

This study has several limitations. As with most human studies of PAE, it is challenging to retrospectively obtain accurate and precise information about PAE. We rigorously screened participant records to confirm PAE all participants, but specific details of dose and timing were often unavailable. PAE is rarely the sole adverse exposure (Astley, 2010), and most participants in this study had additional prenatal or postnatal exposures. Most results remained significant after controlling for these additional exposures, but larger samples are necessary to fully disentangle the effects. Lastly, the elimination of diffusion-weighted volumes can result in an overestimation of diffusion parameters (Chen et al., 2015) however there were no significant differences between groups for number of diffusion-weighted volumes remaining. This study was cross-sectional, which limits the conclusions we can draw about developmental trajectories. Future longitudinal studies are necessary to better characterize the development of brain structure in children with PAE.

## Conclusions

This study demonstrates higher FA and lower MD in white matter in young children with PAE for the first time. These findings provide novel insight into brain development in early childhood of 2-to 7-year-olds with PAE, possibly suggesting premature development. Understanding the structural underpinnings of neurobehavioral symptoms can support approaches for early diagnoses and indicators for early intervention, for young children with PAE.

